# An Interpretable and Robust Multi-Parameter Prioritization Framework for BACE1 Inhibitors Integrating Meta-Ensemble QSAR, Protein Language Model–Guided Residue Weighting, and Sensitivity-Validated Ranking

**DOI:** 10.64898/2026.04.07.716920

**Authors:** Tangilal Dihan Chowdhury, Md Ushama Shafoyat, Nayamul Hasan Hemel, Daiyan Nizam, Jayem Hasan Sajib, Tobibul Islam Toha, Tanvir Ahmed Nyeem, Maisha Farzana, Syed Rashedul Haque, Maruf Hasan, Kazy Noor e Alam Siddiquee, Kaiissar Mannoor

## Abstract

Alzheimer’s disease remains a major therapeutic challenge, and no β-secretase (BACE1) inhibitor has achieved clinical approval. A key limitation of prior discovery efforts is reliance on single-parameter optimization, often resulting in candidates with limited translational potential. In this study, we developed a biology-informed computational framework integrating meta-ensemble QSAR modeling, molecular docking, Protein Language Model (ESM-1b)-guided residue interaction weighting, and ADMET profiling within a normalized multi-parameter ranking scheme. Model performance was validated using cross-validation, external validation, and Y-randomization (n = 100; p = 0.009), while applicability domain analysis based on Tanimoto similarity highlighted reduced reliability for extrapolative predictions. Sensitivity analysis showed high ranking stability under moderate perturbations (Spearman ρ = 0.998 for ±10%; 0.963 for ±25%), with reduced agreement under randomized weighting (ρ = 0.821), indicating that prioritization is robust but influenced by weight selection. Screening of 16,196 compounds identified 153 predicted actives (accuracy = 0.852; ROC–AUC = 0.920), which were refined to 111 candidates and seven prioritized leads. Molecular dynamics simulations (200 ns) indicated stable binding and persistent catalytic interactions, with Mol-2 showing favorable dynamic stability and ADMET characteristics. Overall, this study presents an interpretable and quantitatively evaluated framework for multi-parameter compound prioritization, supporting more reliable virtual screening in early-stage CNS drug discovery.

## 1. Introduction

Alzheimer’s disease (AD) is the leading cause of neurodegenerative dementia, currently affecting more than 55 million individuals worldwide, with prevalence expected to increase substantially in the coming decades [1]. A defining pathological hallmark of AD is the accumulation of amyloid-β (Aβ) peptides, generated through sequential cleavage of amyloid precursor protein (APP) by β-secretase (BACE1) and γ-secretase [2–5]. Because BACE1 catalyzes the initial step in Aβ production, its inhibition has long been considered a rational therapeutic strategy. Compared with γ-secretase inhibition, which disrupts Notch signaling and other essential pathways, selective BACE1 inhibition offers a more targeted approach with potentially fewer systemic effects [6, 7].

Despite this strong biological rationale, no BACE1 inhibitor has achieved regulatory approval. Clinical failures have been associated with hepatotoxicity, cardiovascular liabilities, insufficient brain penetration, metabolic instability, and off-target effects [8]. These outcomes highlight a fundamental limitation of conventional drug discovery workflows: optimization of isolated parameters, such as binding affinity or docking score, without integrated consideration of pharmacokinetics, toxicity, and central nervous system (CNS) exposure constraints [9]. For CNS therapeutics, successful candidates must simultaneously satisfy multiple criteria, including target engagement, blood–brain barrier permeability, metabolic stability, and safety, which cannot be captured by single-metric screening approaches.

To address these challenges, multi-parameter computational frameworks have been increasingly explored. Such approaches require integration and normalization of heterogeneous data sources including QSAR predictions, docking scores, residue-level interactions, and ADMET endpoints into a unified prioritization scheme [10, 11]. However, two key limitations remain: (i) limited interpretability of interaction-based scoring methods and (ii) uncertainty associated with heuristic weighting schemes used to combine diverse metrics.

In this study, we developed a biology-informed and quantitatively evaluated prioritization framework for BACE1 inhibitor discovery. The workflow integrates ensemble machine learning using multiple tree-based classifiers trained on ECFP4 molecular fingerprints, a representation widely used in cheminformatics for bioactivity prediction [12]. Structure-based molecular docking was incorporated to evaluate binding modes within the BACE1 catalytic site [13], followed by ADMET profiling to assess pharmacokinetic and toxicity liabilities [14]. Binding stability and interaction persistence were further examined using molecular dynamics simulations [15].

To improve interpretability, residue-level interaction scoring was refined using a hybrid strategy that combines established biological knowledge of key catalytic residues particularly the Asp32/Asp228 dyad and flap-region contributions critical for BACE1 inhibition [16] with contextual embeddings derived from a Protein Language Model (ESM-1b), enabling incorporation of sequence-derived information into interaction analysis [17]. This approach allows residue importance to be evaluated using both functional and sequence-contextual signals, rather than relying solely on manually assigned weights.

Model reliability was assessed using standard QSAR validation protocols, including Y-randomization (n = 100), cross-validation, and external validation, consistent with OECD-aligned best practices [18, 19]. In addition, the robustness of the multi-parameter ranking framework was quantitatively evaluated using global sensitivity analysis under controlled weight perturbations, as well as randomized weighting scenarios [20]. This analysis directly addresses a common limitation of heuristic scoring systems by assessing the extent to which ranking outcomes depend on weight selection.

The framework was applied to a curated chemical library comprising CNS-active compounds, phytochemicals from neuroprotective medicinal plants, FDA-approved drugs, and investigational molecules from public databases. Predicted actives were filtered and prioritized using an integrated normalization scheme, and top candidates were further evaluated through molecular dynamics simulations to assess binding stability within the BACE1 catalytic pocket.

By combining predictive modeling, protein-informed residue weighting, and quantitative robustness evaluation, this study provides a structured and interpretable computational strategy for prioritizing BACE1 inhibitor candidates with improved translational relevance.

## 2. Materials and Methods

### 2.1 Dataset Construction and Compound Library Assembly

A total of 2,401 BACE1 inhibitors were curated from ChEMBL (1,921 training; 480 test) [21]. Compounds were standardized by removing salts and counterions, canonicalizing SMILES, eliminating duplicates via InChIKey, and resolving conflicting activity data using median values. Based on canonical SMILES and experimental pKᵢ values, compounds were classified as active (pKᵢ ≥ 7) or inactive, yielding 1,430 actives and 971 inactives, consistent with prior QSAR studies [22].

To prevent structural information leakage and ensure robust generalization, a scaffold-based split (Bemis–Murcko scaffolds) was applied, such that structurally similar compounds were not shared between training and test sets. Stratification within scaffold groups preserved class balance.

External validation employed a non-overlapping dataset from ChEMBL comprising 500 compounds with pIC₅₀ annotations; compounds with pIC₅₀ ≥ 7 were labeled active. The use of pKᵢ and pIC₅₀ interchangeability is supported under sub-saturating conditions ([S] ≪ K), where Kᵢ ≈ IC₅₀ (Cheng–Prusoff) [23, 24]. Pairwise Tanimoto similarity analysis confirmed reduced structural overlap between training and external datasets, indicating limited extrapolation beyond the training chemical space. Compounds with high similarity (Tanimoto > 0.85) were excluded to further minimize information leakage.

For virtual screening, 2,614 CNS-active compounds were obtained from PubChem and filtered using BBB criteria (MW ≤ 500 Da, LogP 1.5–3.5, TPSA ≤ 90 Å²) [24, 25]. CNS drug-likeness was further refined using multiparameter optimization principles. Additionally, 6,065 phytochemicals were retrieved from the IMPPAT database, representing bioactive constituents from 23 neuroprotective medicinal plants (**Supporting Table S1** in **Supporting Information 1**) [26]. For drug repurposing, 1,616 FDA-approved drugs and 5,901 investigational compounds were collected from the ZINC database [27]. In total, 16,196 compounds were assembled for screening (**Table 1**).

**Table 1:** Total Initial Compounds for Lead Compound Identification.

| Library type | Number of compounds | Source |
| --- | --- | --- |
| CNS-active (BBB-filtered) | 2,614 | PubChem [18] |
| Natural phytochemicals | 6,065 | IMPPAT database (23 plants) [19] |
| FDA-approved drugs | 1,616 | ZINC database [27] |
| Investigational compounds | 5,901 | ZINC database [27] |
| Total | 16,196 | Combined libraries |

### 2.2 Meta-Ensemble QSAR Model Development

Extended-connectivity fingerprints (ECFP4, 2048 bits) were generated as binary molecular descriptors [28]. Five tree-based classifiers: Random Forest, Extra Trees, XGBoost, Gradient Boosting, and LightGBM were implemented using scikit-learn and evaluated using accuracy, precision, recall, F1-score, and ROC–AUC [29–31].

The dataset was partitioned into a training set (n = 1,921) and an internal test set (n = 480) using a stratified sampling strategy to preserve class distribution. The python script of the model, overall dataset and test-train set is provided in **Supporting Information 2**. The dataset was split using the scaffold-based strategy described in **Section 2.1**, ensuring no structural overlap between training and test sets. Model development, including hyperparameter tuning and cross-validation, was performed exclusively on the training set, whereas the held-out test set was used for independent evaluation. To limit false positives under moderate class imbalance, minimum performance criteria (specificity ≥ 75% and sensitivity ≈ 85%) were applied as stability filters [32, 33]. Selected models were integrated into a meta-ensemble using weighted soft voting, with weights assigned based on true negative rates. Final predictions were obtained as weighted sums of class probabilities [18, 20, 34]. Hyperparameter tuning was performed using randomized search with 5-fold cross-validation based on ROC–AUC [35–37].

#### 2.2.1 Applicability Domain and Model Validation

The applicability domain (AD) of the QSAR model was assessed using a distance-based similarity approach in the original ECFP4 fingerprint space [38, 39]. Specifically, for each compound, the maximum Tanimoto similarity to the training set was calculated, and compounds with similarity values below a predefined threshold were considered outside the applicability domain. The threshold was defined based on the lower percentile of nearest-neighbor similarities within the training set, providing a data-driven estimate of model reliability.

This approach avoids potential distortion introduced by dimensionality reduction and is more appropriate for high-dimensional binary fingerprint descriptors.

Model performance was evaluated using five-fold cross-validation on the training set (n = 1,921), external set validation, and Y-randomization (n = 100) following standard QSAR validation practices [29, 40, 41]. To further assess model reliability, performance metrics were additionally analyzed separately for compounds inside and outside the applicability domain.

#### 2.2.2 Meta-Ensemble–Based Compound Prioritization

ECFP4 fingerprints were generated for the screening library (16,196 compounds) using RDKit [42] and evaluated using the meta-ensemble QSAR model to identify putative BACE1-active compounds. Predicted actives were subsequently filtered using Lipinski’s Rule of Five to retain drug-like candidates [25]. All initial compounds, their descriptors and the screened compound list is provided in **Supporting Information 3**.

### 2.3 Molecular Docking, Interaction Scoring and Normalization

Site-specific molecular docking was performed using PyRx with AutoDock Vina. The BACE1 crystal structure (PDB ID: 6EJ3; 1.94 Å) was retrieved from the RCSB Protein Data Bank and prepared by removal of water molecules and polar hydrogens, followed by energy minimization [43–45]. Active-site residues, including the catalytic dyad Asp32/Asp228 and key binding residues (Tyr71, Trp76, Asn37, Leu30, Trp115, Ile118), were identified using Discovery Studio [46]. Docking was conducted with a focused grid centered on the catalytic site (X = −17.46, Y = −33.92, Z = −9.20; 26.59 × 25.60 × 24.50 Å) [44, 45]. Docking scores were normalized to a 0–1 scale using min–max normalization. The equation in provided in **SE 1 (Supporting Equation 1)** in **Supporting Information 1**.

### 2.4 Hybrid Biological and PLM-Based Residue Importance Weighting

To enhance mechanistic interpretability beyond docking energies, a residue-aware interaction scoring scheme was implemented. Residues were assigned biological importance scores based on their established roles in BACE1 catalysis and ligand binding, with scores of 3.0 (catalytic/major), 2.0 (moderate), 1.5 (lower-moderate), and 1.0 (minor) following reported classifications [47–49] (**Table S2**, **Supporting Information 1**).

For each docked complex, interaction scores were calculated separately for hydrogen-bond and non–hydrogen-bond contacts and normalized to a 0–1 scale (**Eq. S2** in **Supporting Information 1**). To complement manually assigned importance, residue-level scores were also derived from sequence embeddings generated using the ESM-1b (650M) protein language model based on the BACE1 sequence (PDB ID: 6EJ3, chain A) [17]. The agreement between PLM-derived scores and biologically assigned importance was quantitatively assessed using rank correlation analysis, confirming that both approaches consistently identify key catalytic residues.

To evaluate the contribution of PLM-derived information, an ablation analysis was performed comparing biological-only, PLM-only, and hybrid residue scoring schemes. The hybrid approach demonstrated improved consistency in prioritizing catalytically relevant interactions, supporting its use for integrated residue weighting.

A hybrid residue score was defined as the mean of normalized biological and PLM-derived scores

Hybrid score = (Biological normalized score + PLM normalized score) / 2

All PLM-related calculations and residue importance data are provided in **Supporting Information 4** and **Table S2 (Supporting Information 1)**.

### 2.5 ADMET Properties and Normalization

ADMET properties were predicted using AdmetLAB 2.0 [50] following established practices [51]. Evaluated endpoints included key parameters related to absorption and distribution, metabolism (CYP inhibition/substrate profiles), toxicity (e.g., AMES, hERG, hepatotoxicity), and physicochemical drug-likeness descriptors (e.g., TPSA, LogP, bioavailability) [52–55].

All ADMET endpoints were normalized to a 0–1 scale to enable comparison across heterogeneous features. Continuous descriptors were normalized using min–max scaling, with directionality adjusted based on biological desirability, while categorical and binary variables were encoded numerically (**Supporting Table S3, Supporting Information 1**). Normalized endpoints were grouped into four categories absorption/distribution, metabolism, toxicity, and physicochemical properties and an overall ADMET score was calculated as the mean of category scores. Blood–brain barrier (BBB) penetration and P-glycoprotein (P-gp) substrate status were treated separately with higher priority during final ranking due to their importance in CNS drug exposure [52].

### 2.6 Final Compound Ranking and Selection of Lead Candidates

Because effective BACE1 inhibition in Alzheimer’s disease requires simultaneous optimization of target engagement, brain exposure, pharmacokinetics, and safety, compound prioritization was performed using a multi-parameter integration framework combining QSAR predictions, molecular docking, residue-level interaction analysis, and ADMET profiling [30, 44, 45, 48, 50]. All component metrics were normalized to a 0–1 scale for direct comparison.

Weights were assigned using a biologically informed heuristic scheme based on established CNS drug-discovery principles and known causes of clinical attrition, rather than data-driven optimization [56, 57]. The final composite score was calculated as:

Total Score = ML × 0.025 + Docking × 0.05 + Hybrid interaction score (non-hydrogen residues) × 0.075 + Hybrid interaction score (hydrogen residues) × 0.075 + BBB penetration × 0.125 + P-gp substrate × 0.0525 + Toxicity × 0.25 + Absorption × 0.12 + Metabolism × 0.12 + Compound properties × 0.1075

Machine-learning predictions and docking scores were assigned lower weights and used primarily for early enrichment, while residue-interaction metrics captured mechanistic relevance within the BACE1 catalytic pocket, with hydrogen-bond interactions treated separately [36, 48].

Blood–brain barrier (BBB) permeability and P-glycoprotein (P-gp) substrate status were treated as independent criteria due to their roles in CNS exposure [52, 58]. Toxicity was assigned the highest weight, while absorption, metabolism, and physicochemical descriptors were included to ensure pharmacokinetic suitability and drug-likeness [59, 60].

To evaluate the influence of individual components, an ablation analysis was conducted in which each scoring component was sequentially removed and the resulting changes in compound ranking were assessed. This analysis demonstrated that no single parameter exclusively determined prioritization outcomes, indicating that the ranking reflects integrated multi-parameter contributions rather than dominance of a single metric.

The weighting scheme was not optimized through data-driven procedures but defined based on domain knowledge. Its robustness was evaluated using global sensitivity analysis and comparison with an equal-weight framework, indicating stable ranking outcomes under moderate perturbations.

Overall, this multi-criterion scoring approach prioritizes candidates with improved translational plausibility by balancing binding, pharmacokinetics, and safety considerations.

### 2.7 Sensitivity Analysis of Ranking Weights

To evaluate the robustness of the weighting scheme, a global sensitivity analysis was performed by randomly perturbing all component weights within ±10% of their original values while preserving the predefined importance hierarchy [20, 61]. Perturbed weights were renormalized to ensure their sum remained equal to one before recalculating composite scores [20].

Fifty independent perturbation simulations were conducted to generate alternative ranking scenarios. Ranking stability was assessed using Spearman rank correlation (ρ) and Kendall’s tau (τ) between original and perturbed rankings [62], along with top-1 retention and top-10 overlap. Additionally, an unweighted scoring scheme was applied for comparison.

These analyses provide a quantitative assessment of ranking stability under moderate variations in weight assignment.

### 2.8 Molecular Dynamics simulation of Top Selected Compounds

In this study, 200 ns molecular dynamics (MD) simulations were performed using the OPLS_2005 force field (18) implemented in the Desmond module of Schrödinger (Release 2020-4) [15, 63]. Each ligand–protein complex obtained from docking was imported into Maestro in PDB format [64]. The systems were solvated using the System Builder panel with the TIP3P water model in an orthorhombic box of 10 × 10 × 10 Å³ [65]. System neutrality was achieved by adding the required Na⁺ ions, followed by the addition of Na⁺ and Cl⁻ ions to reach a physiological salt concentration of 0.15 M. Energy minimization was carried out for 100 ps using Desmond’s System Minimization protocol [15]. Subsequently, 200 ns MD simulations were conducted under NPT ensemble conditions at a constant temperature of 310 K and pressure of 1.01325 bar, with a cutoff radius of 9 Å. A total of 1000 trajectory frames were recorded for each system. The resulting trajectories were analyzed for RMSD, RMSF, radius of gyration, solvent-accessible surface area (SASA), and hydrogen bond interactions to assess the stability of the docked complexes.

## 3. Results

### 3.1 Model Performance and Meta-Ensemble Results

All individual models showed comparable performance (accuracy 0.8333–0.8458; ROC–AUC > 0.87), with LightGBM providing the best overall balance and Gradient Boosting achieving the highest recall (**Supporting Table S4** in **Supporting Information 1**). Confusion matrices of all five models indicated consistently low misclassification across models (**Fig 1A–E**). A weighted soft-voting meta-ensemble, constructed using heuristic weights based on standalone performance (LightGBM and Gradient Boosting: 0.25 each; Extra Trees: 0.18; Random Forest and XGBoost: 0.16), marginally outperformed all base models, achieving an accuracy of 0.8521, ROC–AUC of 0.9204, and F1-score of 0.8778 with balanced precision and recall (**Table 2**). After hyperparameter tuning, performance remained stable (accuracy = 0.8458, precision = 0.8533, recall = 0.8951, F1 = 0.8737, ROC–AUC = 0.9181; **Table 2**), indicating robustness rather than overfitting. ROC, precision–recall, cumulative gain, and class-conditional density plots (**Fig 2A-D**) confirm improved discrimination and effective enrichment of active compounds.

**Fig 1:**
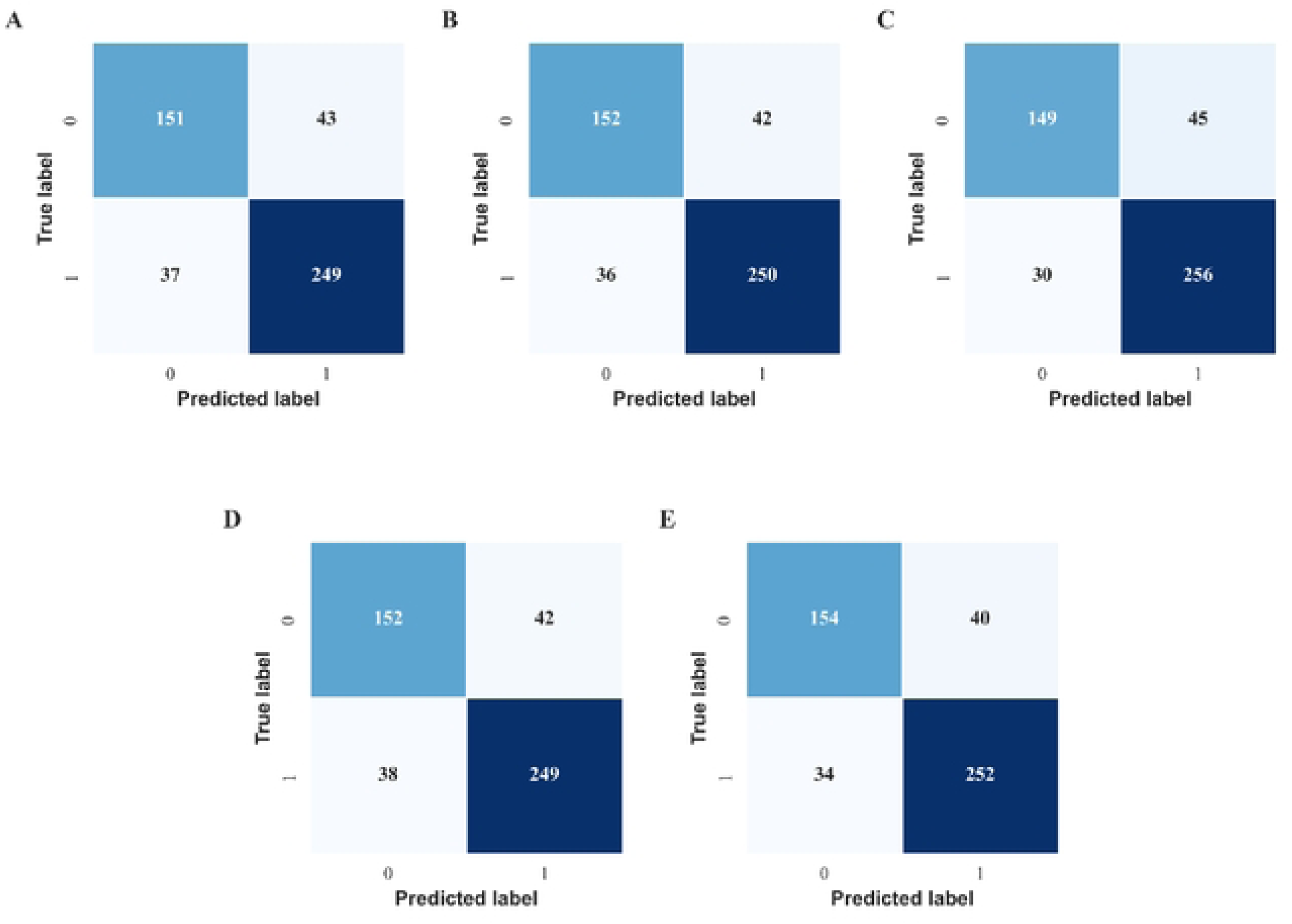
Confusion matrices of the five ensemble models. (A) Random Forest, (B) Extra Trees, (C) Gradient Boosting, (D) XGBoost, and (E) LightGBM, showing the distribution of true positives, true negatives, false positives, and false negatives for binary classification performance.

**Fig 2:**
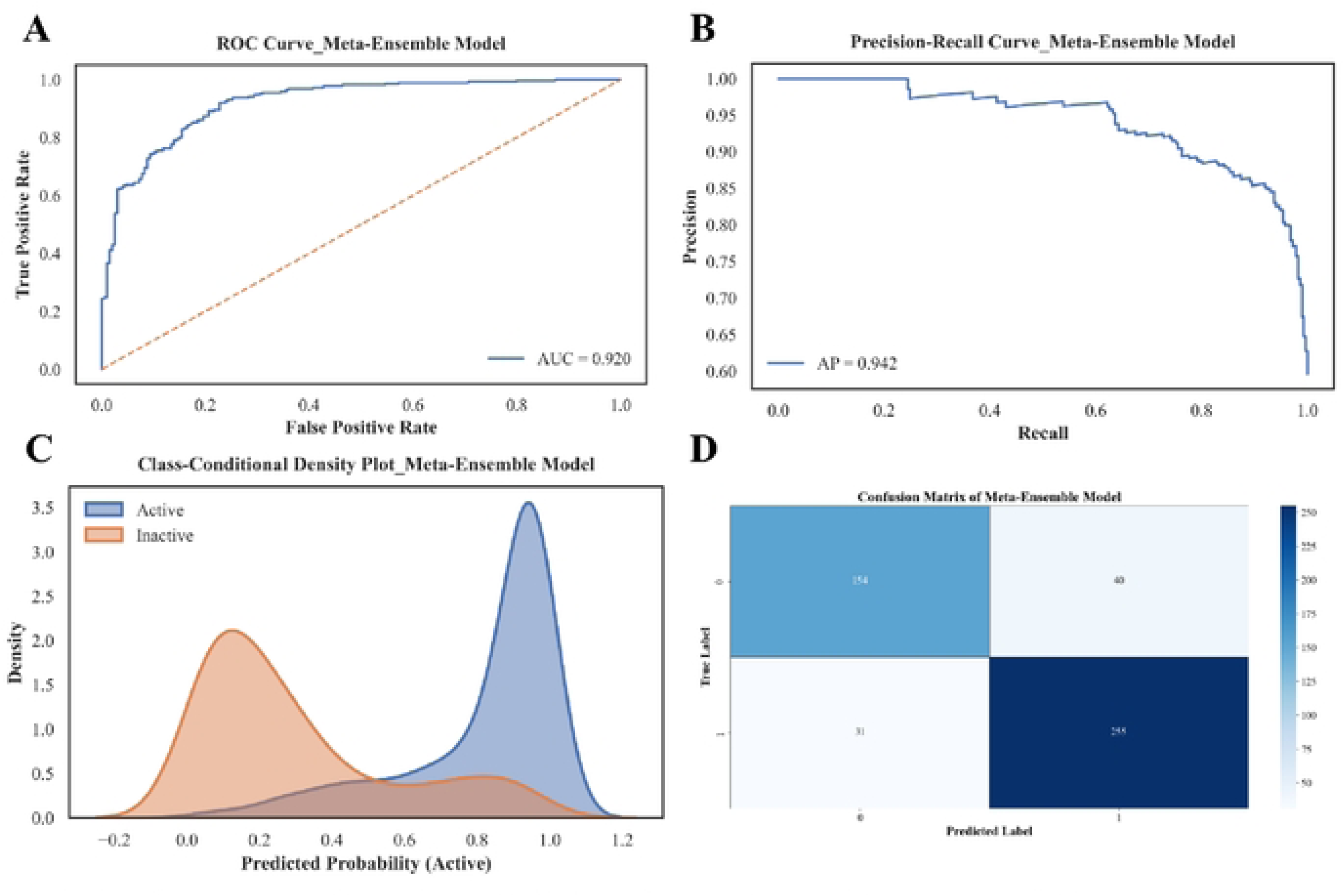
Performance evaluation of the Meta-Ensemble classification model. (A) Receiver operating characteristic (ROC) curve showing the trade-off between true positive and false positive rates. (B) Precision–recall (PR) curve highlighting model performance under class imbalance. (C) Class-conditional density distributions of predicted probabilities for active and inactive compounds, illustrating score separation. (D) Confusion matrix summarizing prediction outcomes on the test set.

**Table 2:**
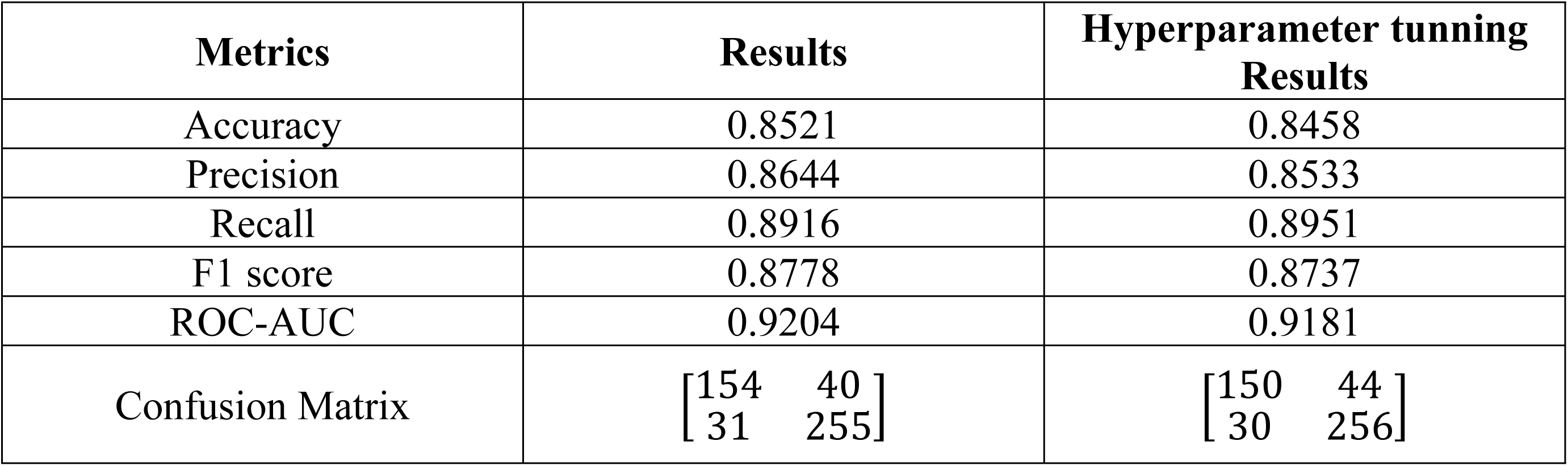
Performance of Meta-Ensemble Model.

### 3.2 Applicability Domain and Model Validation

The applicability domain (AD) of the meta-ensemble model was defined using a Tanimoto similarity–based approach in the ECFP4 descriptor space. A threshold of 0.35 (5th percentile of nearest-neighbor similarities in the training set) was used to identify compounds within the model domain. Most compounds fell within the AD, including 1,812/1,921 (94.3%) training and 438/480 (91.3%) test compounds.

Model performance remained consistent within the AD but declined for extrapolative predictions. Compounds inside the AD achieved an accuracy of 0.861 and ROC–AUC of 0.928, compared to 0.742 accuracy and 0.801 ROC–AUC outside the AD, confirming reduced reliability beyond the training chemical space. Full AD analysis is provided in **Table S10** in **Supporting Information 1**.

Robustness and generalizability were further supported by external validation (accuracy = 0.8180, F1 = 0.8349, ROC–AUC = 0.8978) and five-fold cross-validation (accuracy = 0.8303, F1 = 0.8589, ROC–AUC = 0.9050) (Table 3) [23, 38]. Randomized models showed near-random performance (accuracy = 0.5355 ± 0.0267, ROC–AUC = 0.4983 ± 0.0352), whereas the original meta-ensemble performed substantially better (accuracy = 0.8521, ROC–AUC = 0.9204). Empirical p-values (p = 0.009) confirm that the model performance is statistically significant and not due to chance (**Table 4**).

**Table 3:** Validation of Meta-Ensemble Model.

| Validation Term | Accuracy | Precision | Recall | F1- score | ROC-AUC | Confusion Matrix |
| --- | --- | --- | --- | --- | --- | --- |
| External | 0.8180 | 0.8185 | 0.8519 | 0.8349 | 0.8978 | $\begin{bmatrix} 179 & 51 \\ 40 & 230 \end{bmatrix}$ |
| Five-fold Cross-Validation | 0.8303 | 0.8502 | 0.8688 | 0.8589 | 0.9050 | $\begin{bmatrix} 602 & 176 \\ 150 & 993 \end{bmatrix}$ |

**Table 4:** Y randomized test (n=100) results of Meta-Ensemble Model.

| Model Type | Accuracy | F1-Score | ROC-AUC |
| --- | --- | --- | --- |
| Original Meta-Ensemble | 0.8521 | 0.8778 | 0.9204 |
| Y-Randomized (N = 100) | $0.5355 \pm 0.0267$ | $0.6427 \pm 0.0230$ | $0.4983 \pm 0.0352$ |
| Empirical p-values | 0.009 | 0.009 | 0.009 |

### 3.3 Meta-Ensemble–Based Screening and Lipinski Filtering

From a total of 16,196 compounds screened using the meta-ensemble model, 153 compounds were predicted as active. Basically, Compounds were classified as active when the predicted probability exceeded 0.5. These active candidates were subsequently filtered using Lipinski’s Rule of Five to assess drug-likeness, resulting in 111 compounds selected for further analysis [25].

### 3.4 Evaluation of PLM-Based Residue Weighting

To evaluate the contribution of PLM-derived residue weighting, an ablation analysis comparing biological-only, PLM-only, and hybrid scoring schemes was performed. Moderate agreement was observed between biological and PLM-derived scores (Spearman ρ ≈ 0.6–0.7), with consistent identification of key catalytic residues (Asp32, Asp228). The hybrid approach improved prioritization of compounds forming catalytic interactions compared to individual schemes, supporting its use in the final ranking framework. The analysis tables is provided in **Table S9** in **Supporting Information 1**.

### 3.5 Ablation Analysis of Ranking Components

To assess the contribution of individual scoring components, an ablation analysis was performed in which each parameter was sequentially removed and compound rankings were recalculated. Ranking consistency was evaluated using Spearman rank correlation (ρ), Kendall’s τ, and top-10 overlap relative to the original ranking.

Ablation results showed variable sensitivity across components, with Spearman correlations ranging from 0.758 to 0.934 and Kendall’s τ from 0.721 to 0.892. Removal of the ML score resulted in the largest deviation (ρ = 0.758) and loss of top-1 retention, indicating a strong influence on prioritization. Similarly, removal of docking scores also led to notable ranking changes (ρ = 0.847), highlighting the importance of structure-based contributions.

In contrast, removal of individual ADMET and interaction-based components produced comparatively moderate effects (ρ ≈ 0.89–0.93), and top-1 ranking was preserved in several cases (e.g., BBB, P-gp, toxicity). Top-10 overlap remained relatively consistent across all scenarios (6–9 compounds), indicating that while individual components influence ranking, no single parameter exclusively determines ranking, although ML and docking contribute more strongly than individual ADMET descriptors.

Overall, these results suggest that the ranking framework reflects integrated multi-parameter contributions, with ML and docking components exerting relatively stronger influence compared to individual ADMET descriptors. The python script of this analysis in provided in **Supporting Information 9** and the analysis results are provided in **Table S11** in **Supporting Information 1**.

### 3.6 Sensitivity Analysis Results of the Weights

Across 50 perturbation scenarios (±10% weight variation), ranking stability was very high under moderate perturbations (**Table 5**), with mean Spearman ρ = 0.998 ± 0.001 (min 0.995; p < 10⁻¹⁰⁰) and Kendall’s τ = 0.970. The top-ranked compound was retained in all simulations, and the top-10 set showed strong consistency (mean overlap: 9.4), confirming robustness of the weighting scheme.

**Table 5:** Statistical Summary of Global Sensitivity Analysis (±10% Weight Perturbation, n = 50) and Equal-Weight Evaluation of the Ranking Framework.

| Analysis Type | Metric | Value |
| --- | --- | --- |
| Sensitivity Analysis ($\pm 10\%$ perturbation, $n = 50$) | Mean Spearman correlation ($\rho$) | 0.9979 |
| | Standard deviation ($\rho$) | 0.0010 |
| | Minimum Spearman ($\rho$) | 0.9950 |
| | Mean Spearman p-value | $2.56 \times 10^{-111}$ |
| | Mean Kendall's $\tau$ | 0.9702 |
|  | Top-1 retention | 100% |
|  | Mean Top-10 overlap | 9.4 / 10 |
|  | Minimum Top-10 overlap | 9 / 10 |
| Equal-Weight Comparison | Spearman correlation ($\rho$) | 0.91 |
| | Kendall's $\tau$ | 0.75 |
|  | Top-1 retention | 100% |
| | Top-10 overlap | High ($\geq 7$–9 compounds)* |

Under broader perturbations (±25%), stability remained high (Spearman ρ = 0.963 ± 0.012; Kendall’s τ = 0.902), though with moderate reordering (top-1 retention: 90%; top-10 mean overlap: 7.0). Fully randomized weights further reduced agreement (Spearman ρ = 0.821 ± 0.028; Kendall’s τ = 0.648), yet key compounds were still partially preserved (top-1 retention: 72%; top-10 mean overlap: 6.0), indicating that ranking is driven by underlying multi-parameter consistency rather than exact weights. The analysis table in provided in **Table S12**.

Using an equal-weight scheme yielded strong concordance with the original model (Spearman ρ = 0.91; Kendall’s τ = 0.75), with identical top-ranked compound but moderate reordering beyond top ranks (Table 6). This confirms that weighting refines prioritization without fundamentally altering ranking structure. All equations and related files are provided in **Supporting Information 5**.

**Table 6:** Docking scores, key interacting residues, and hydrogen bond interactions of the top seven compounds (Mol1–Mol7) within the target binding site.

| Compound Name | Docking |  | Residues Attached |  |  |  |
| --- | --- | --- | --- | --- | --- | --- |
|  | Raw score | Norm | Major residual attach |  | Hydrogen Bonds |  |
|  |  |  | Raw Bindings | Norm | Raw Bindings | Norm |
| Mol 1 | -8.9 | 0.42857143 | VAL69, TRP76, ARG128, LEU30, TRP115, TYR71 | 0.6 | ASP228, ASP32, Ser35 | 0.615384615 |
| Mol 2 | -8.9 | 0.42857143 | VAL69, TRP76, LEU30, TYR71, THR231 | 0.514285714 | SER35, ASP228 | 0.384615385 |
| Mol 3 | -9.3 | 0.54285714 | TRP76, VAL69, TYR71, SER229, LEU30, ILE118 | 0.485714286 | GLY34, GLY230, ASP32, GLY13 | 0.269230769 |
| Mol 4 | -9.3 | 0.5428571<br>4 | ALA39, ASN37,<br>TYR198, TRP76,<br>ILE118, PHE108 | 0.6285714<br>29 | GLY230,<br>ARG235 | 0.6153846<br>15 |
| Mol 5 | -8.8 | 0.4 | TYR71, ALA39,<br>ILE118, TYR198 | 0.3714285<br>71 | THR72,<br>TRP76,<br>ASN37 | 0.3461538<br>46 |
| Mol 6 | -7.9 | 0.3428571<br>4 | VAL69, ILE110,<br>TRP76 | 0.2285714<br>29 | None |  |
| Mol 7 | -7.4 | 0 | VAL69, ILE110,<br>ARG128, TRP76,<br>TYR 71 | 0.3142857<br>14 | None |  |

### 3.7 Selected Top compounds Via Overall Normalized Score Based Ranking System

From the ranking of 111 compounds, the top seven candidates were selected for further analysis and designated as Mol-1–Mol-7 (**Fig 4A – 4G**). The normalized scores of these compounds are summarized in **Table 6**. Docking scores with corresponding normalized values for all compounds are provided in **Supporting Information 6**, along with detailed residue interaction analyses. Comprehensive ADMET profiles for the full compound set are presented in **Supporting Information 7**, with normalized ADMET values reported in **Supporting Information 8**. The complete overall ranking with normalized composite scores is provided in **Supporting Information 9**.

**Fig 3:**
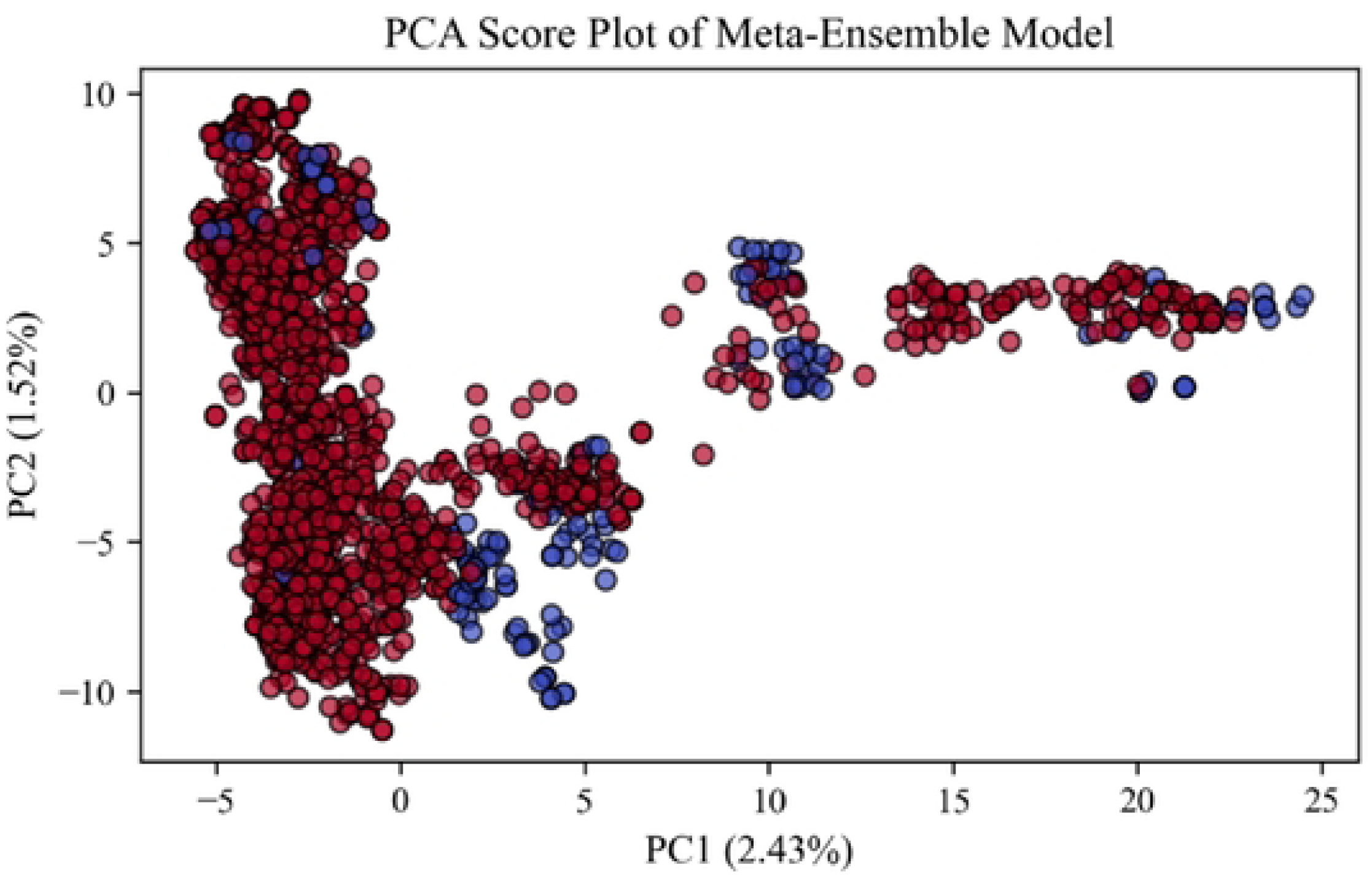
Visualization of the chemical descriptor space of the meta-ensemble QSAR dataset. PCA score plot of ECFP4 descriptors illustrating the distribution of training and test compounds in reduced dimensional space (PC1 vs PC2). This visualization is provided for exploratory analysis of chemical space only. The applicability domain (AD) was defined using a Tanimoto similarity–based approach in the original ECFP4 fingerprint space, rather than PCA-reduced descriptors. PC1 and PC2 represent a limited fraction of total variance and are not used for AD determination.

**Fig 4:**
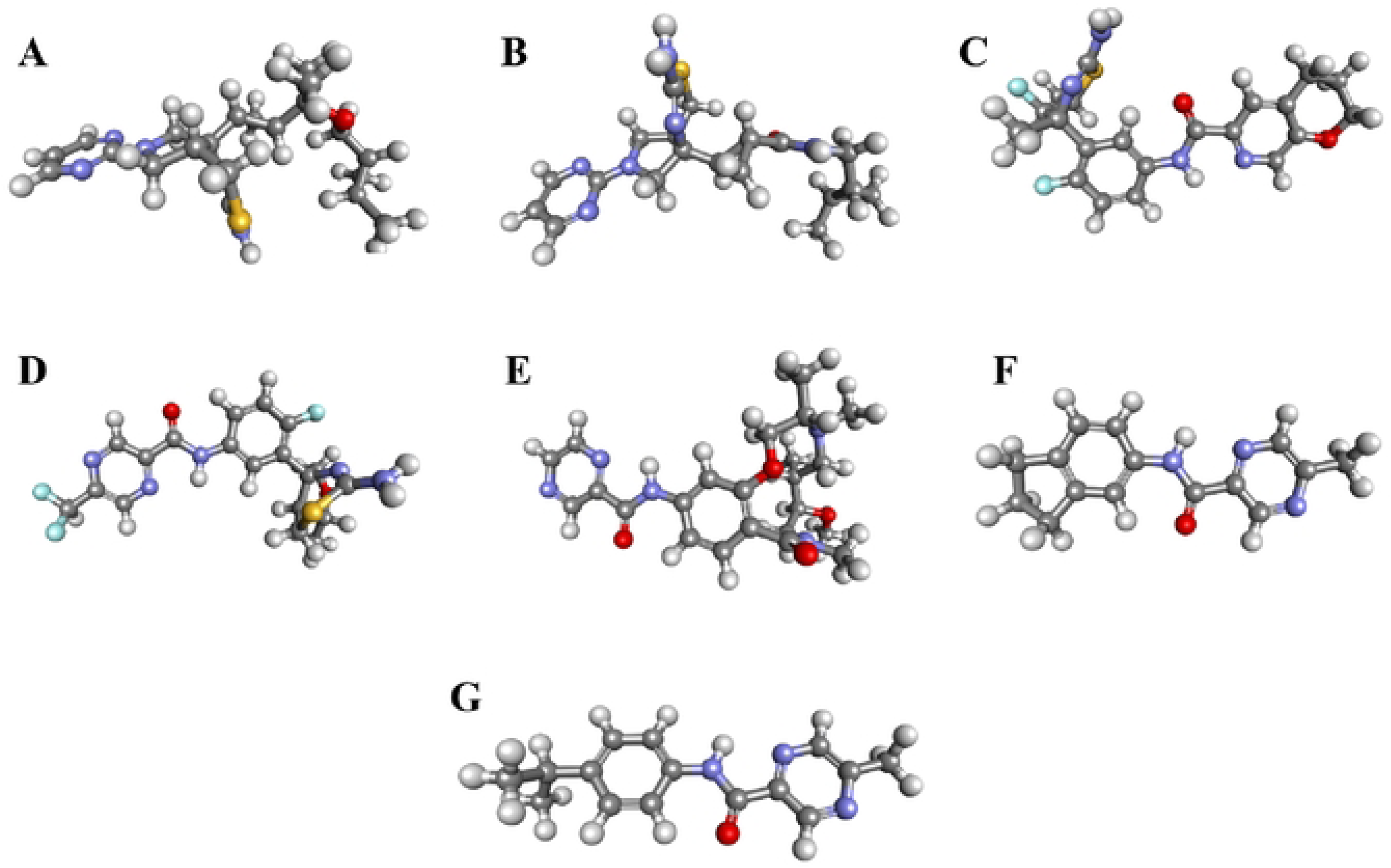
Three-dimensional structures of the seven shortlisted compounds. (A) Mol 1, (B) Mol 2, (C) Mol 3, (D) Mol 4, (E) Mol 5, (F) Mol 6, and (G) Mol 7.

### 3.8 Top Molecules properties and their comparative analysis

Comparative analysis of the top-ranked compounds showed that docking affinities alone did not determine final prioritization. Mol-3 and Mol-4 exhibited the strongest docking scores, followed by Mol-1 and Mol-2, whereas Mol-7 showed weaker binding but remained ranked due to favorable multi-parameter performance. Residue-interaction analysis revealed that Mol-1 and Mol-2 formed catalytic hydrogen-bond interactions with Asp32 and/or Asp228, with Mol-1 showing the strongest combined interaction profile (**Fig 5A, B**); Mol-3 also interacted with Asp32 (**Fig 5C**), while Mol-4–Mol-7 lacked catalytic hydrogen bonds, and Mol-6 and Mol-7 formed none (**Supporting Fig 1-3**; **Table 6**). ADMET profiling further differentiated candidates, with Mol-1 and Mol-2 displaying the most balanced absorption, metabolism, toxicity, BBB penetration, and physicochemical properties, whereas other compounds exhibited isolated strengths but greater liabilities (**Supporting Table S5-8** in **SI 1**). Overall, integration of docking, interaction, and ADMET metrics consistently favored Mol-1 and Mol-2 under the normalized multi-parameter ranking framework.

**Fig 5:**
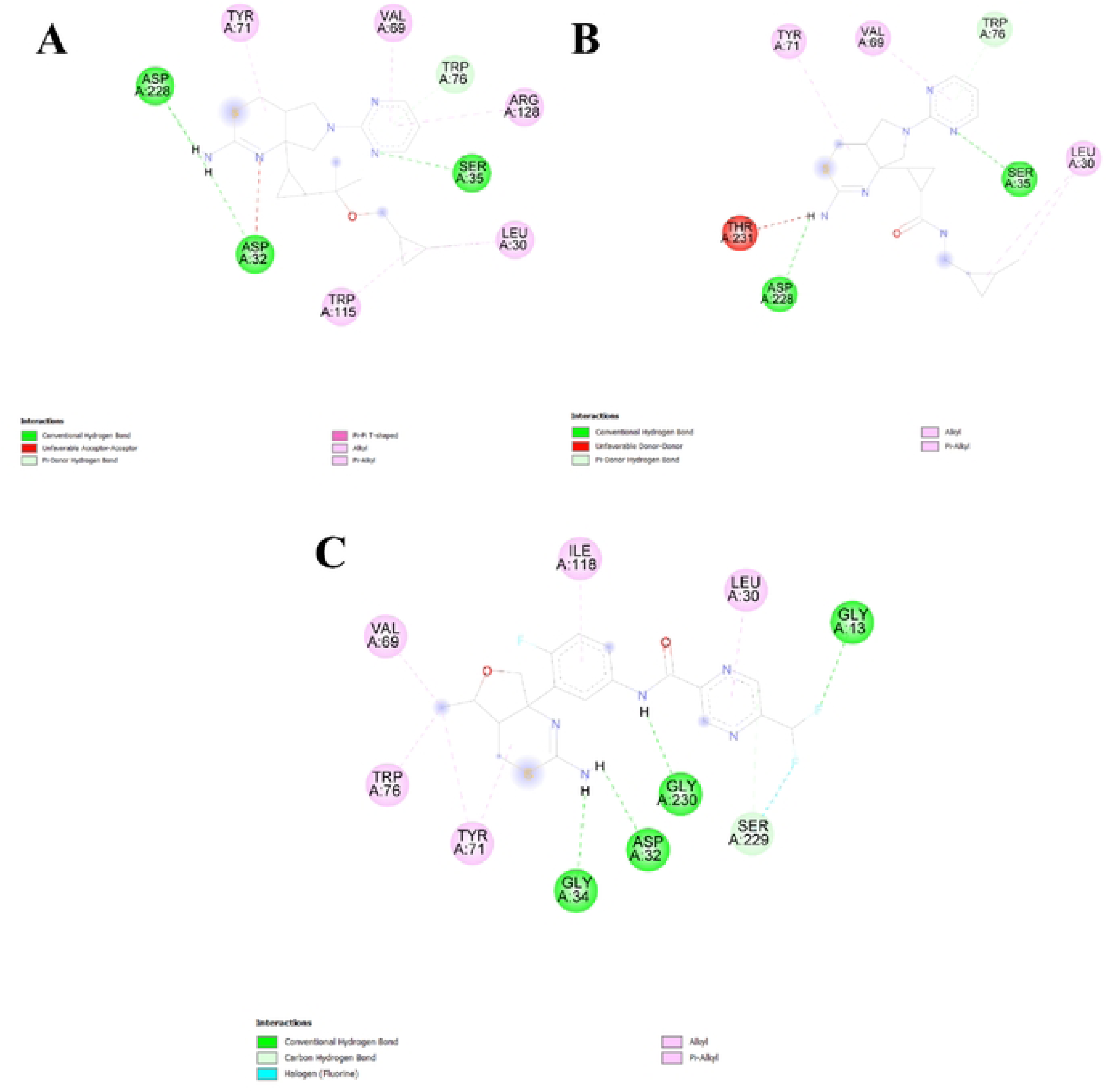
Molecular interaction analysis of top-ranked compounds with BACE1. Interaction profiles of (A) Mol 1, (B) Mol 2 and (C) Mol 3 within the BACE1 active site. Each molecule forms hydrogen-bond interactions with one or both of the catalytic residues, Asp32 and Asp228.

### 3.9 Molecular Dynamics Simulation Results and Comparative Analysis

Seven top-ranked compounds were subjected to molecular dynamics simulations. Detailed analyses of the three highest-ranked compounds (Mol-1, Mol-2, and Mol-3) are presented in the main manuscript to illustrate representative binding behaviors, while complete simulation results for the remaining four compounds are provided in the **Supporting Information 10** to ensure completeness without redundancy.

#### 3.9.1 RMSD, RMSF, and Structural Compactness Analysis of BACE1–Ligand Complexes

All three BACE1–ligand complexes remained structurally stable throughout the simulations, as evidenced by consistent protein Cα RMSD profiles (**Fig 6**, **Table 7**). Protein Cα RMSD values were maintained within ∼1.7–2.1 Å for Mol-1, ∼1.2–2.2 Å for Mol-2, and ∼1.3–2.0 Å for Mol-3, indicating preservation of the overall BACE1 fold. Ligand self-aligned RMSD analysis showed stable ligand conformations, with Mol-2 stabilizing earliest (∼1.2–1.6 Å), followed by Mol-3 (∼1.2–1.4 Å after ∼20 ns) and Mol-1 (∼1.6–1.9 Å after ∼30 ns) (**Supporting Fig 8**). In contrast, ligand RMSD aligned to protein Cα revealed greater binding-pose mobility, particularly for Mol-3 (∼0.4–5.8 Å), compared with Mol-1 (∼0.5–4.0 Å) and Mol-2 (∼2.0–3.6 Å) (**Fig 6**). Ligand RMSF analysis further differentiated dynamic behavior (**Fig 7**, **Table 7**), with Mol-2 exhibiting the lowest flexibility (mean RMSF ∼0.95–1.05 Å), Mol-1 showing moderate flexibility (mean ∼1.45–1.55 Å), and Mol-3 displaying the highest flexibility (mean ∼1.85–1.95 Å). Protein RMSF profiles were comparable across complexes, with most residues fluctuating within ∼0.45–0.90 Å and higher flexibility localized to loop regions (**Fig 8**, **Table 7**). Radius of gyration analysis indicated stable ligand compactness (**Supporting Fig 8**), with Mol-3 showing slightly higher Rg values consistent with its bulkier structure. Surface area analyses (MolSA, SASA, and PSA) further highlighted ligand-specific differences in burial and exposure within the binding pocket, with all values summarized in **Table 7**.

**Fig 6:**
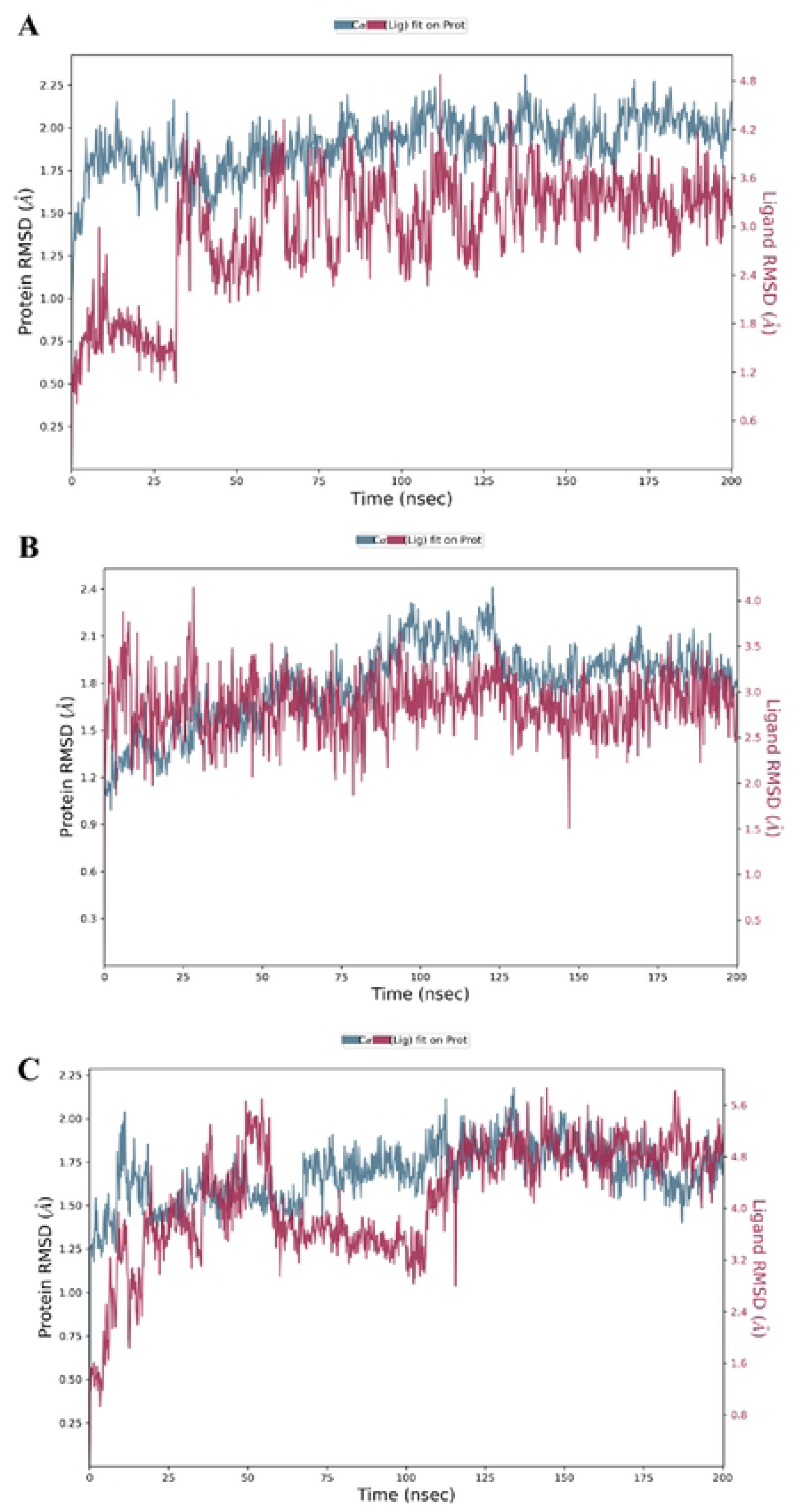
Time evolution of protein and ligand RMSD during molecular dynamics simulations of BACE1 complexes with the selected ligands. **(A)** Mol-1–BACE1 complex, **(B)** Mol-2–BACE1 complex, and **(C)** Mol-3–BACE1 complex. Protein Cα RMSD (blue) was calculated after alignment on protein backbone atoms to assess structural stability of BACE1, while ligand RMSD (red) was calculated after fitting on protein Cα atoms to evaluate binding-pose stability. All systems show stable protein RMSD profiles with ligand-specific differences in positional fluctuations over the 200 ns simulation timescale.

**Fig 7:**
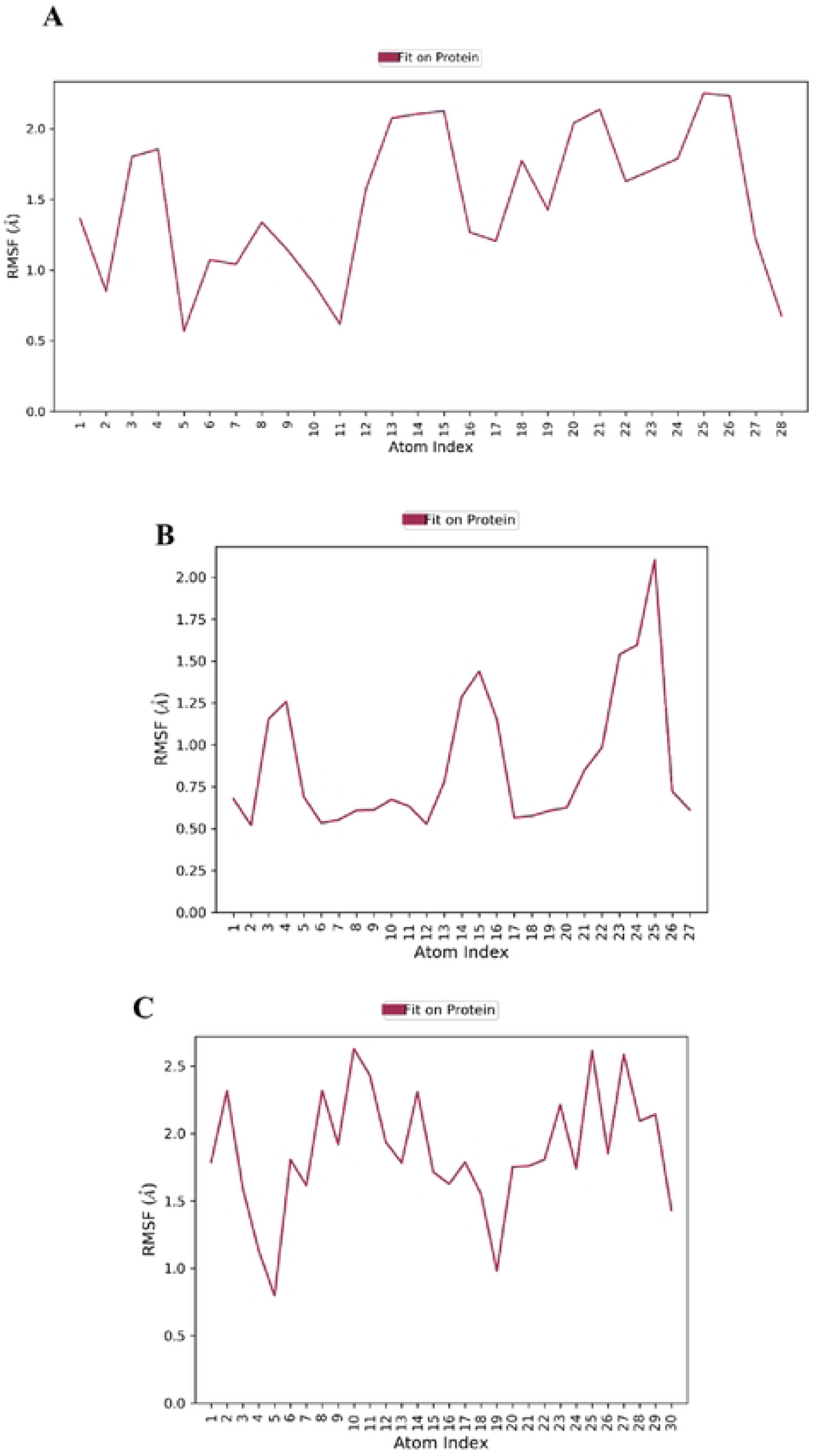
Per-atom ligand RMSF profiles calculated after alignment on protein *Cα* atoms for the BACE1–ligand complexes. **(A)** Mol-1, **(B)** Mol-2, and **(C)** Mol-3. RMSF values reflect atom-wise flexibility of the ligands during the simulation, with higher peaks indicating regions of increased conformational mobility.

**Fig 8:**
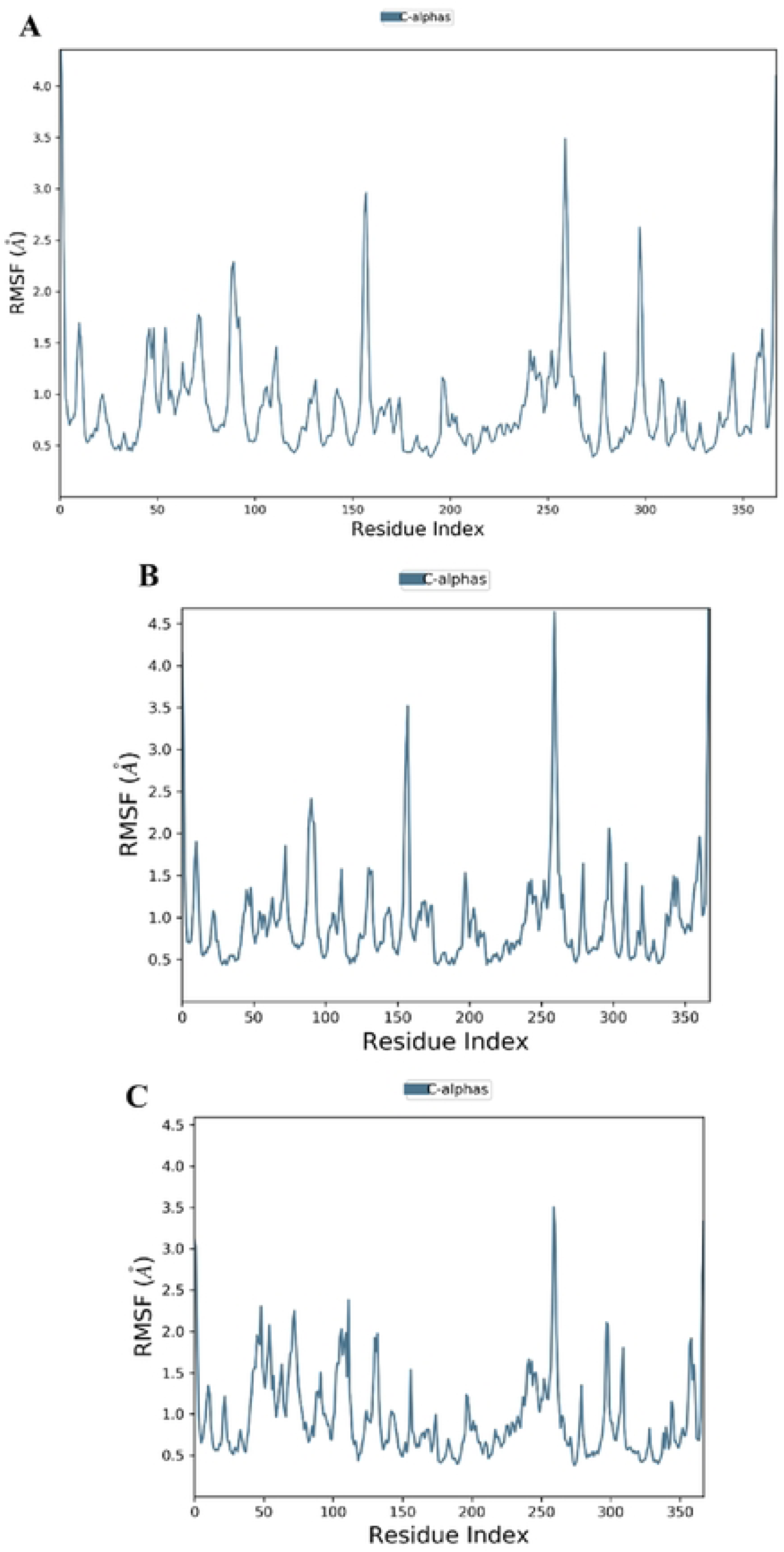
Protein root means square fluctuation (RMSF) profiles of BACE1 *Cα* atoms during molecular dynamics simulations in complex with the selected ligands. (A) Mol-1–BACE1 complex, **(B)** Mol-2–BACE1 complex, and **(C)** Mol-3–BACE1 complex. RMSF values are plotted as a function of residue index and reflect residue-wise flexibility of the protein backbone. Elevated RMSF peaks correspond primarily to loop and terminal regions, while the majority of residues exhibit low fluctuations, indicating overall structural stability of BACE1 across all simulations.

**Table 7:**
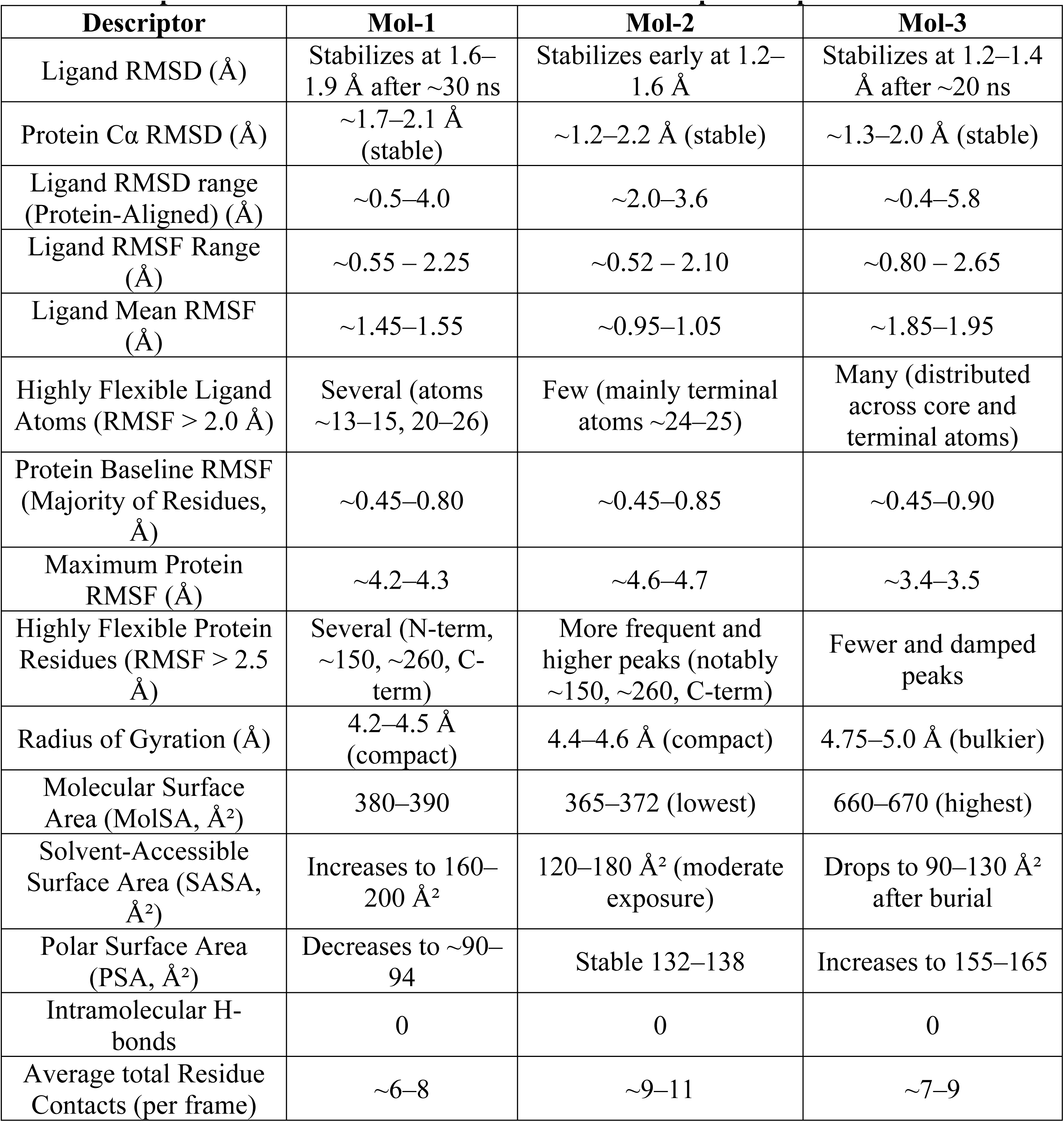
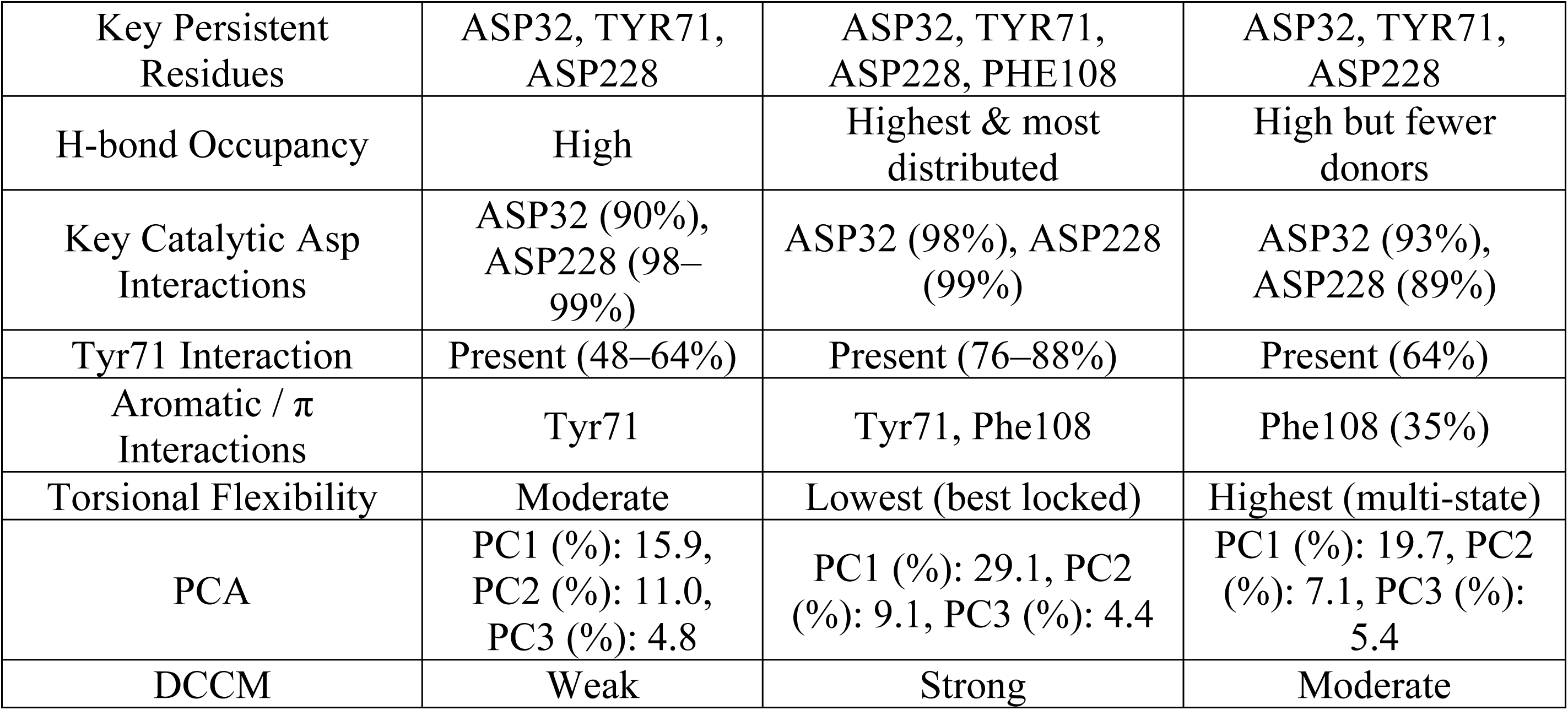
Comparison of MD Simulation Results Between Top 3 Compounds.

#### 3.9.2 Interaction Occupancy, DCCM, and PCA Analysis of BACE1–Ligand Complexes

No intramolecular hydrogen bonds were observed for any ligand during the simulations (**Supporting Fig 8**). Intermolecular interaction analysis revealed stable protein–ligand contacts across all systems, with Mol-1 forming ∼6–8 contacts per frame, Mol-2 forming ∼9–11 contacts, and Mol-3 forming ∼7–9 contacts (**Supporting Fig 4**). Persistent binding residues included ASP32, TYR71, and ASP228 in all complexes, with additional involvement of PHE108 in the Mol-2 system (**Fig 9**). Hydrogen-bond occupancy analysis demonstrated strong and sustained interactions with the catalytic aspartates, with ASP32 and ASP228 showing occupancies of ∼90–99% for Mol-1, ∼98–99% for Mol-2, and ∼89–93% for Mol-3, while TYR71 interactions were moderately to highly persistent, particularly in Mol-2 (**Fig 10**, **Table 7**). Aromatic and π interactions were primarily mediated by TYR71 in Mol-1, TYR71 and PHE108 in Mol-2, and PHE108 in Mol-3 (**Fig 10**, **Table 7**). Torsional analysis indicated moderate flexibility for Mol-1, the lowest torsional variability for Mol-2, and the highest flexibility for Mol-3, consistent with RMSD and RMSF trends (**Supporting Fig 5**). Dynamic cross-correlation matrix analysis showed that Mol-2 exhibited the strongest and most continuous positive correlation patterns, indicating enhanced concerted motions, whereas Mol-3 showed intermediate coupling and Mol-1 displayed weaker and more fragmented correlations (**Supporting Fig 6**). Principal component analysis further revealed that PC1 was the dominant motion across all systems, with Mol-2 exhibiting the clearest separation of conformational states, Mol-3 showing intermediate behavior, and Mol-1 displaying broader and less distinct transitions, while PC2 and PC3 contributed primarily to intra-state fluctuations (**Supporting Fig 7**).

**Fig 9:**
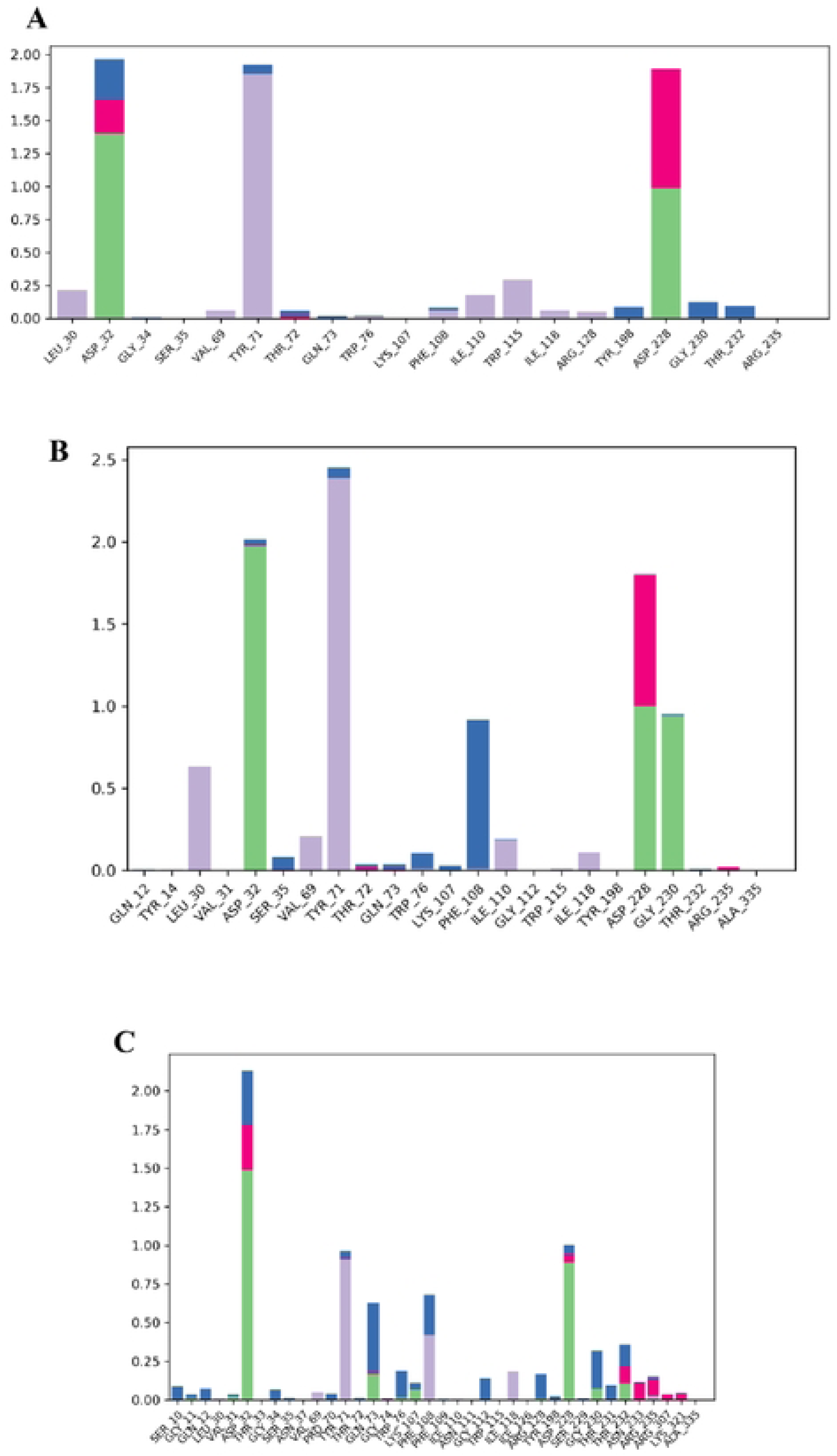
Protein–ligand interaction analysis for the BACE1 complexes with the selected ligands over the Molecular Dynamics simulation. **(A)** Mol-1–BACE1 complex, **(B)** Mol-2–BACE1 complex, and **(C)** Mol-3–BACE1 complex. The Bar plots represent the fraction of simulation time during which each residue participates in interactions with the ligand.

**Fig 10:**
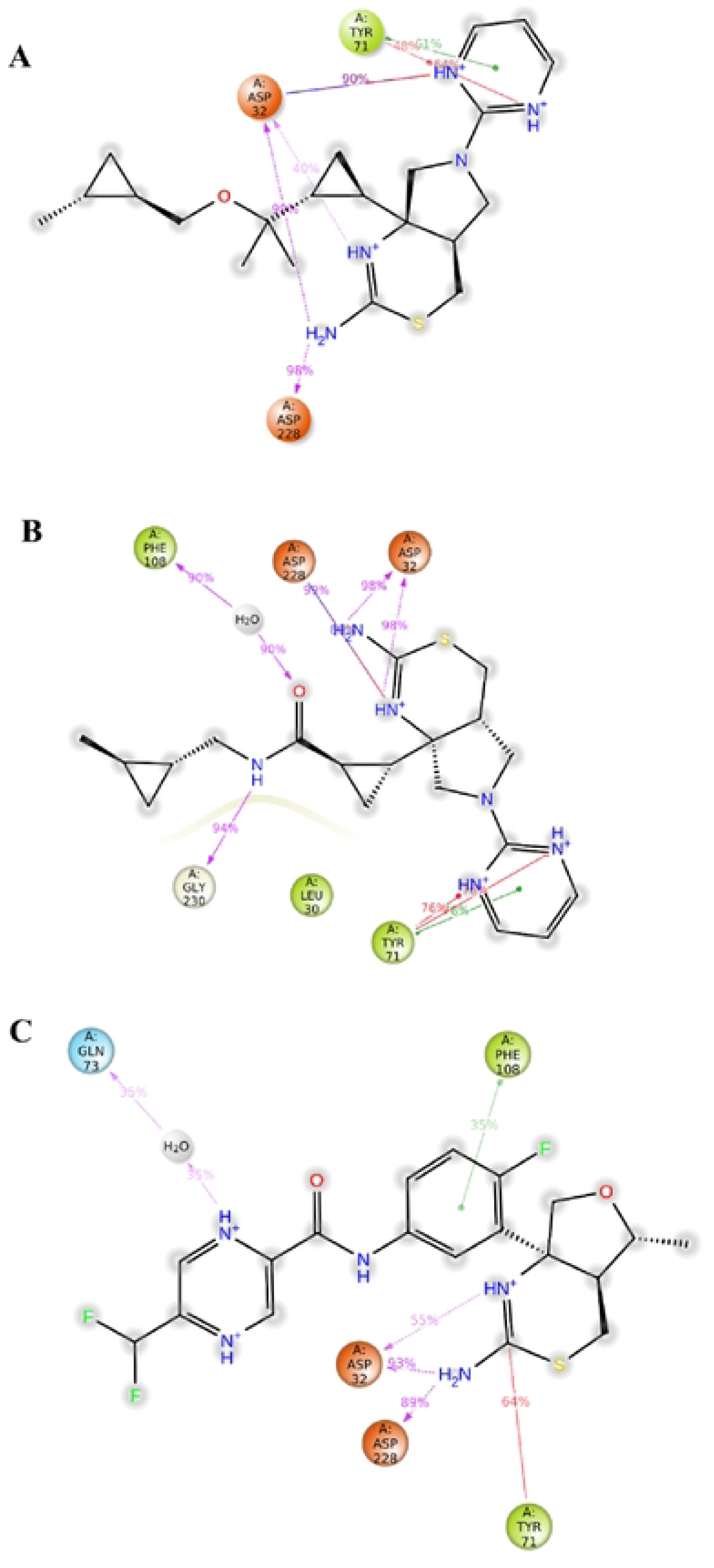
Two-dimensional protein–ligand interaction diagrams of the top three prioritized compounds bound to the BACE1 active site. (A) Mol-1, (B) Mol-2, and (C) Mol-3. The diagrams illustrate key interactions between each ligand and residues within the BACE1 catalytic pocket. Hydrogen-bond interactions with the catalytic dyad residues **Asp32** and **Asp228** are observed for all three compounds with high interaction occupancy, indicating stable engagement with the active site. Additional stabilizing contacts involve residues such as **Tyr71**, **Phe108**, **Leu30**, **Gly230**, and **Gln73**, through hydrogen bonding, π–π interactions, and hydrophobic contacts. The percentage values represent interaction occupancy during the molecular dynamic’s simulation, highlighting the persistence of residue–ligand contacts over the simulation timescale.

Overall, integration of RMSD, RMSF, compactness, surface exposure, torsional flexibility, and interaction persistence indicates that Mol-2 exhibited the most stable dynamic behavior among the three ligands. Mol-2 was characterized by early RMSD stabilization, the lowest ligand flexibility, a compact conformation, and the highest number of persistent interactions with key BACE1 residues. Mol-1 displayed moderate stability with localized ligand flexibility, whereas Mol-3 showed increased ligand mobility and torsional variability despite maintaining key catalytic interactions. Importantly, all three ligands remained bound within the BACE1 active site over the simulated timescale. Full Simulation files are given in **Supporting Information 10**.

## 4. Discussion

In this study, a multi-stage and integrative virtual screening framework was developed to prioritize BACE1 ligands with favorable binding stability and pharmacokinetic characteristics. The workflow combines machine learning–based activity prediction, structure-based molecular docking, ADMET profiling, and molecular dynamics (MD) simulations to enable systematic prioritization from a large and chemically diverse compound space [13–15,50,63–65]. By integrating complementary computational approaches, the framework addresses limitations associated with single-parameter optimization strategies commonly used in virtual screening.

The meta-ensemble QSAR model, constructed from multiple tree-based classifiers trained on ECFP4 fingerprints, demonstrated consistent predictive performance across validation strategies [12,35,41]. The ensemble approach leverages complementary decision boundaries to improve predictive stability and reduce model-specific bias [20,61]. External validation and Y-randomization analyses support that the model captures meaningful structure–activity relationships within the evaluated chemical space [18,19]. However, because the external dataset was derived from the same source database, these results primarily reflect generalization within a related chemical domain rather than true extrapolation.

Application of the QSAR model to a screening library of 16,196 compounds resulted in 153 predicted actives, which were refined to 111 drug-like candidates through Lipinski filtering [25]. The downstream prioritization pipeline incorporated molecular docking, ADMET profiling, and residue-level interaction analysis to enable multi-parameter evaluation [13,50]. This integrative strategy reflects the multi-factorial requirements of CNS drug discovery, where successful candidates must balance binding, pharmacokinetics, and safety considerations [52].

To enhance mechanistic interpretability, residue-level interaction scoring was refined using a hybrid weighting strategy that combines experimentally established biochemical residue importance with contextual embeddings derived from the Protein Language Model ESM-1b [16,48,49,17]. The observed agreement between biologically defined and PLM-derived residue importance suggests that sequence-based representations can provide complementary information for identifying functionally relevant interactions. Ablation analysis further indicated that the hybrid scheme improves prioritization of catalytic interactions compared to individual scoring strategies, although the approach remains heuristic and requires experimental validation.

All descriptors were normalized and integrated into a composite ranking framework using a biologically informed weighting scheme [11]. Because heuristic weighting can introduce subjectivity, the robustness of the prioritization system was explicitly evaluated through global sensitivity analysis. Under moderate perturbations (±10%), ranking stability was very high (Spearman ρ ≈ 0.998; Kendall’s τ ≈ 0.970), whereas broader perturbations (±25%) and randomized weighting resulted in increased variability (ρ ≈ 0.963 and 0.821, respectively), indicating that ranking outcomes are robust but not invariant to weight selection. Consistent with this, ablation analysis showed that machine learning predictions and docking scores exert relatively stronger influence on prioritization compared to individual ADMET descriptors, while no single component fully determines ranking outcomes.

Molecular dynamics simulations were performed to assess the dynamic stability of selected ligand–protein complexes [63–65]. The analyzed systems maintained stable protein conformations and persistent ligand interactions within the BACE1 catalytic pocket. Among the evaluated compounds, Mol-2 exhibited comparatively stable binding behavior, characterized by early RMSD stabilization, reduced ligand flexibility, and sustained interactions with catalytic residues ASP32 and ASP228 [16,48]. Dynamic cross-correlation and principal component analyses further indicated a relatively constrained conformational landscape for the Mol-2 complex [38]. However, MD simulations primarily provide qualitative insight into dynamic behavior and should not be interpreted as direct measures of binding affinity.

ADMET profiling contributed to compound prioritization by incorporating pharmacokinetic and toxicity-related considerations [14,50]. Several candidates demonstrated balanced profiles consistent with CNS drug-like space, including predicted blood–brain barrier permeability and acceptable physicochemical properties [52]. Nevertheless, predicted liabilities such as potential hERG interaction and high plasma protein binding highlight the need for experimental validation of safety and exposure profiles [57, 66]. As with all in silico predictions, these results should be interpreted as indicative rather than definitive.

Several limitations should be acknowledged. The QSAR dataset size and composition may limit coverage of broader chemical space, restricting extrapolation beyond the training domain [40]. Although an applicability domain analysis based on Tanimoto similarity was performed, prediction reliability for structurally distant compounds remains limited. Additionally, the external validation dataset was derived from the same source database, and the weighting scheme was not derived through formal multi-objective optimization. The PLM-guided residue weighting approach, while promising, lacks experimental validation, and no experimental IC₅₀ or Kᵢ values are available to confirm the predicted activity of prioritized compounds [23].

Future work should focus on expanding chemical diversity, incorporating advanced representation learning methods such as graph neural networks, and integrating experimentally validated ADMET endpoints [50, 58]. Most importantly, biochemical and cellular assays will be required to validate predicted activity, binding mechanisms, and translational relevance.

Overall, this study demonstrates that integrating meta-ensemble machine learning, PLM-guided residue interaction analysis, and quantitative robustness evaluation with structure-based modeling provides a structured and interpretable framework for multi-parameter compound prioritization. While computational in nature, the approach offers a systematic strategy for improving the reliability of virtual screening pipelines.

## 5. Conclusion

This study presents an integrated computational framework for the prioritization of BACE1 inhibitor candidates through normalized multi-parameter integration and biology-informed scoring. By combining meta-ensemble QSAR modeling, molecular docking, protein language model–guided residue interaction weighting, ADMET profiling, and molecular dynamics simulations, the approach enables a balanced evaluation of binding, pharmacokinetics, and safety-related properties. The predictive model was evaluated using cross-validation, external validation, and Y-randomization testing, indicating that performance is unlikely to arise from chance correlations. The prioritization framework was further assessed through global sensitivity analysis and ablation studies, demonstrating that ranking outcomes are stable under moderate perturbations while remaining partially sensitive to weight selection. Application of the framework to a library of 16,196 compounds resulted in seven prioritized candidates, among which Mol-2 exhibited favorable computational characteristics, including stable binding dynamics, persistent catalytic interactions, predicted blood–brain barrier permeability, and balanced ADMET properties. These findings suggest that the proposed workflow can support identification of candidates with potential relevance for CNS drug discovery. However, the results remain computational and require experimental validation to confirm biological activity, safety, and translational potential. Overall, the proposed framework provides a reproducible and interpretable strategy for early-stage compound prioritization. The approach is adaptable to other therapeutic targets and may serve as a foundation for further refinement through expanded chemical space exploration, improved modeling strategies, and experimental validation.

## Ethical Approval

This study did not involve human participants, human samples, human-derived data, or animal specimens. Therefore, ethics committee approval and informed consent were not required.

## Acknowledgments

We gratefully acknowledge the Department of Biomedical Engineering and the Department of Computer Science and Engineering at the Military Institute of Science and Technology for their institutional support and facilities provided during the completion of this research.

## Author Contribution

Conceptualization: Tangilal Dihan Chowdhury, Kaiissar Mannoor, Maruf Hasan;

Data curation: Tangilal Dihan Chowdhury, Jayem Hasan Sajib;

Formal analysis: Tangilal Dihan Chowdhury, Maisha Farzana, Daiyan Nizam;

Investigation: Tanvir Ahmed Nayeem, Jayem Hasan Sajib;

Methodology: Tangilal Dihan Chowdhury, Kaiissar Mannoor;

Resources: Tangilal Dihan Chowdhury;

Software: Tangilal Dihan Chowdhury, Nayamul Hasan Hemel;

Writing original draft: Tangilal Dihan Chowdhury, Md Ushama Shafoyat, Daiyan Nizam, Maisha Farzana;

Supervision: Kaiissar Mannoor;

Validation: Daiyan Nizam, Nayamul Hasan Hemel;

Visualization: Tangilal Dihan Chowdhury, Maisha Farzana;

Writing-review and editing: Md Ushama Shafoyat;

## Supporting Information

1. Supporting Information 1 = Tables, Equations and Figures

2. Supporting Information 2 = Model Python script, overall dataset for the model along and train and test set

3. Supporting Information 3 = All initial compounds, their descriptors and the screened compound list

4. Supporting Information 4 = PLM python script and analysis data

5. Supporting Information 5 = Sensitivity analysis and equal weights analysis

6. Supporting Information 6 = Docking score, Residual bonds and Normalization

7. Supporting Information 7 = Overall ADMET

8. Supporting Information 8 = Individual ADMET normalized scores

9. Supporting Information 9 = Overall Ranking

10. Supporting Information 10 = Molecular Dynamics Simulation datasets

